# Development of social behaviour in young zebrafish

**DOI:** 10.1101/017863

**Authors:** Elena Dreosti, Gonçalo Lopes, Adam R. Kampff, Stephen W. Wilson

## Abstract

Adult zebrafish are robustly social animals whereas larva is not. We designed an assay to determine at what stage of development zebrafish begin to interact with and prefer other fish. One week old zebrafish do not show significant social preference whereas most 3 weeks old zebrafish strongly prefer to remain in a compartment where they can view conspecifics. However, for some individuals, the presence of conspecifics drives avoidance instead of attraction. Social preference is dependent on vision and requires viewing fish of a similar age/size. In addition, over the same 1–3 weeks period larval zebrafish increasingly tend to coordinate their movements, a simple form of social interaction. Finally, social preference and coupled interactions are differentially modified by an NMDAR antagonist and acute exposure to ethanol, both of which are known to alter social behavior in adult zebrafish.

## Introduction

Human infants exhibit social behaviours from birth(Xiao et al., 2014). Throughout life these innate social drives provide the substrate for learning more complex forms of human interaction. Disruptions to early social behaviour may impair the development of normal adult sociality, and may contribute to disorders such as autism (Banerjee et al., 2014). Since the neural circuitry that underlies human innate social behaviour is established *in utero*, very little is understood about its normal and pathological development, anatomy, and function.

Early developing social behaviours, such as the preference to observe and mimic conspecifics, are common to many other mammals (Ferrari et al., 2006) and non-mammalian vertebrates (Mooney, 2014; Engeszer et al., 2004; 2007). Animal models are much more amenable to detailed investigation and share many of the same anatomical and functional neural systems that underlie innate social behaviour in humans (O’Connell and Hofmann, 2012; 2011). Consequently, we sought a model system for which neural circuits can be assessed throughout development, and for which social behaviour is an important component of the organism’s behavioural repertoire (Oliveira, 2013).

Zebrafish adults are social animals(Oliveira, 2013), exhibiting a range of group (shoaling and schooling) (Krause et al., 2000; Miller and Gerlai, 2012; Green et al., 2012), conspecific directed aggression (Jones and Norton, 2015), mating (Engeszer et al., 2008) and other behaviours(Arganda et al., 2012). Larval zebrafish, however, do not exhibit the overt shoaling and schooling behaviours that are readily apparent in adults. In order to shoal fish must prefer to approach and remain near conspecifics. There is some evidence that this preference might develop as early as one week (Hinz et al., 2013) whereas shoaling appears only in early flexion larvae (~ 13–15 days post fertilization (dpf, 6 mm length) (Engeszer et al., 2007). Schooling, however, requires the group to move in a polarized and coordinated manner and has thus far only been described in adults(Miller and Gerlai, 2012).

Social behaviour encompasses more than simply preferring to be near members of the same species. For instance, individuals may coordinate their behaviour with other members of the same social group. Such coordination is obvious in the case of schooling fish, where individuals align their body orientation and synchronize their movements, but it is also present in social mammals. For example, humans will unconsciously coordinate a diverse range of behaviours, such as yawning, eye blinks and posture (Richardson et al., 2007; Sebanz et al., 2006), and this is thought to provide a foundation for more elaborate forms of social communication and cooperation.

Here we set out to investigate early social interactions in zebrafish and to determine if the establishment of preference for the presence of conspecifics is contemporaneous with individuals beginning to coordinate their behaviour. We have designed a novel social preference/interaction assay for zebrafish larvae that continuously monitors, with high temporal resolution, the detailed behaviour of individuals freely choosing to observe or avoid a counterpart. This assay demonstrates that social behaviours develop gradually and are robust by three weeks post-fertilisation. We have also used the assay to characterize the effects of substances known to influence the social behaviour of adults.

## Materials and Methods

*Zebrafish husbandry.* AB strain zebrafish *(Danio rerio)* to be tested were bred, raised and housed in the same environment. All fish were obtained from natural spawning and housed in groups of roughly 50 fish, and kept at 14h light/10h dark cycle. Fish were fed two times per day from 4 dpf with dry food diet from SAFE Diets (particle size 50–100) and twice with salt water rotifer (Brachionus Plicatilis) until 10 dpf; then twice a day with dry food diet (particle size 100–200), and with a combination of salt water rotifer and brine shrimp (Artemia salina) until 15 days; finally twice a day with dry food diet (particle size 20300) and with brine shrimp until used in the experiments. All the fish run in the behavioural assay were fed in the morning. All test fish were paired with age-matched siblings as the social cue. The experiments described were approved by local ethical committee and the UK Home Office.

*Behavioural assay.* Experiments were performed in a custom-built behavioural setup (Fig. 1A) that was assembled from structural framing (Misumi, DE) and optomechanics (Thorlabs, USA). The videography system comprised a high-speed camera (Flea3, PointGrey, CA), infrared LED backlight (Advanced Illumination, USA, 880 nm), infrared filter (R70, Hoya, JP), and a vari-focal lens (Fujinon, JP). Fish were imaged in a custom-built behavioural arena that was fabricated with a laser-cutter from 5 mm thick opaque white acrylic, sealed with silicone, and with glass window partitions; the multi-chamber design is shown in Fig. 1A. The dimensions of a single behavioural arena was 4 cm × 3.2 cm. The viewing chamber that contained a single or multiple SC fish was 1.5 cm square, and the width of the passage between the two arms of the arena was 6 mm. The water level height was 5 mm, the temperature of the water was around 25’C, pH was 7.0, and the conductivity was 445 µS.The arena was supported on a transparent base covered on one side with diffusive gel paper (Rosco Cinegel, USA). It was illuminated with visible light by homogenously projecting a white rectangle, via a 45° infrared cold mirror positioned between the chamber and IR illuminator, onto the base of the assay using a laser light projector (Microvision, ShowwX+, USA). For all experiments, the entire behavioural apparatus was enclosed in a light-tight enclosure, and for the dark experiments, the visible background illumination was removed.

**Figure 1.** Social preference is robust in three week old zebrafish. **A,** Schematic of the behavioural setup (top). Infrared light homogeneously illuminates the behavioural arenas. Schematic of a single choice chamber (bottom left) with an observer (test) fish and multiple conspecifics as the social cue (SC). The blue lines are clear glass windows. Single frame from high-speed video recording of an experiment with three week old fish (bottom right). **B,** Examples of tracking of a one week old (top) and three week old (bottom) fish, in the absence (left) and presence (right) of the SC. The blue and red portions of the movement tracks are used to calculate the social preference index (SPI, indicated below). **C,** Schematic depicting body orientation of the observer test fish relative to the SC chamber (inset - top left). Polar histograms, averaged across all tested fish, of body orientations of the observer fish when within the SC side of the chamber. From one to three weeks a preference emerges for the observer fish to view the SC with either the left −45°) or right (+45°) eye. Thin lines indicate two standard errors from the mean (SEM) (one week: n=143, two weeks: n=151, three weeks: n=181). **D,** Histograms of all SPIs during acclimation (left column) and SC (right column) periods across different developmental stages (one week (− dpf); two weeks (13–15 dpf), three weeks (20–22 dpf). A range of positive and negative preferences are observed. For presentation clarity, red bars (SPI > 0.5) highlight strong preference for the SC, while blue bars (SPI < −0.5) highlight strong aversion for the SC (zero is marked with a dashed vertical line). **E,** Histogram of all SPIs when a single conspecific served as the SC across different developmental stages. Numbers in brackets indicate Mean SPI.

*Acquisition software:* Fish in six individual chambers were contemporaneously tracked in real-time using custom written workflows in Bonsai, an open-source C# data stream processing framework(Lopes et al., 2015). For each chamber, the image was cropped, background subtracted, thresholded, and the centroid found, characterized (position, orientation, and heading), and stored in a CSV file. The video was also saved with H.264 compression for subsequent offline analysis (https://www.dreo-sci.com/resources/).

*Data analysis:* Social Preference Index (SPI) was calculated by subtracting the number of frames in which the fish was located within the region near the conspecific SC (area highlighted by the red tracking in Fig. 1B) from the number of frames spent in the equivalent region on the opposite side of the chamber (blue tracking in Fig. 1B). This difference was then divided by the total number of frames recorded in the two analysis compartments during the experiment, resulting in a value varying between − 1 and 1. [“SPI = (SC frames – Non SC frames)/ Total frames”]. The SPI during the acclimation period, for which there is no SC, was computed with reference to the randomized side of the chamber on which the SC would be added in the subsequent experimental phase.

The compressed video from each experiment could be repeatedly re-analysed using custom written bulk-processing routines in Python (https://www.dreo-sci.com/resources/). A motion trajectory for each fish was computed by first segmenting the binary particle for each fish from the background and then measuring the change in pixel values, for that particle, from one frame to the next. This resultant frame-by-frame segmented motion value provided a very stable time series for identifying the peaks of individual bouts and then testing for interaction between the observer and SC fish.

*Statistical analysis:* The SPI distributions for many Ac vs SC conditions (one week old, MK-801, EtOH, etc.) are normally distributed, however some conditions (in which the value approaches the SPI bound of +/− 1), are clearly non-normal. We therefore used the same non-parametric statistic test for all comparisons in the study: a Wilcoxon signed-rank test of paired samples.

*Drug treatments:* MK-801: 100 mM stock solution was prepared by dissolving MK- 801 (M107; Sigma-Aldrich) in 100% DMSO (D2650; Sigma-Aldrich) and stored at − 20°C. The drug was administered for 1 h prior the experiments by diluting the stock solution in fish water in order to obtain a working concentration of 100 μM. Zebrafish were washed with fish water before placing them in the chamber for recordings.

For ethanol experiments, low (0.125%) or a high ethanol (0.5%) concentrations were obtained by diluting ethanol in fish water. Fish were exposed with one of the two ethanol concentrations for 1 h prior to, and during experiments.

## Results

### Fish develop strong social preference and interactions by three weeks of age

We designed a behavioural chamber in which zebrafish fish could swim freely between two arms, but in only one they could view conspecific siblings through a glass partition. Six such chambers were simultaneously monitored with an infrared high-speed camera and automated tracking software recorded the behaviour (position and orientation) of the observer fish (Fig. 1A, Movie 1). Following 15 minutes in the chamber without conspecifics (acclimation (Ac) period), a single or three conspecifics were added to one of the adjacent compartments, randomly selected, and the behaviour of the observer fish was monitored for an additional 15 minutes (social cue (SC) period). There was no bias between compartment arms in the acclimation phase for fish at any age, nor if the fish were monitored for a further 15 minutes following the acclimation phase without adding the SC (Supp. Figure 1E-F).

Three week old zebrafish consistently showed a very strong bias to remain in the arm of the chamber adjacent to the SC (Fig. 1B, compare Supp. Movie 2 (one week) to Supp. Movie 3 (three week)). To quantify the tendency for each tested fish to spend time in one or the other arm of the chamber, we defined a social preference index (SPI) (see Materials and Methods). A positive SPI indicates a preference for the chamber arm with the SC and a negative SPI indicates an aversion for the SC. The SPI was computed for all tested one, two and three week old fish with and without the presence of multiple (Fig. 1D) or a single conspecific (Fig 1E). One week old larvae exhibited a very weak, but significant, preference for an SC arm (Fig. 1D, top; Ac vs SC p=0.006) containing multiple conspecifics, however, this preference bias was absent when only a single fish was placed in the SC arm (Fig. 1E, top; Ac vs SC p=0.9). In contrast, the SPI of two week old larvae was strongly shifted towards positive values when viewing both multiple (Fig. 1D, middle; Ac vs SC, p=4.4*10^−10^) and single conspecifics (Fig. 1E, middle; Ac vs SC, p=1.4*10^−11^, Supp. Fig. 1D). By three weeks, SC preference strengthened further, with many values close to 1, reflecting the strong bias of some observer fish to remain almost entirely on the side of the conspecifics (multiple: Fig. 1D, bottom; Ac vs SC, p=2.0*10^−13^, single: Fig. 1E, bottom; Ac vs SC,p=1.4*10^−15^).

A small minority of three week old fish had strong negative SPIs (Fig 1D, bottom). These fish exhibited an aversive response to the conspecifics, preferring to stay in the chamber away from the SC (Supp. Fig. 1B and Supp. Movie 4). Such aversive behaviour was rarely observed in younger fish suggesting that, as for positive social interaction, social aversion also increases throughout development.

The behaviour of the three week old zebrafish when viewing the SC consisted of alternating body orientation such that the left or right eye directly viewed the SC compartment (Supp. Movie 3). This behaviour was quantified in a histogram of all orientations of the fish body axis while in the SC arm of the chamber (Fig. 1C, Supp. Fig. 1C). A gradual transition from primarily orienting along the axes of the chamber (cardinal directions: 0°, +/− 90°, and 180°) to orienting for visual observation (+/− 45°) occurred over the first three weeks of development. No strong bias for observing with either the left or right eye was found in this assay (Sovrano and Andrew, 2006).

We next set out to investigate what sensory cues contribute to the displayed social preference.

### Social preference requires visual observation of conspecifics with a similar age

Although visual stimulation seemed the most likely source of the preference for the SC, it was possible that some olfactory or tactile cues may pass between the chambers of the observer and SC fish. Consequently, we compared preference behaviour for three week old fish tested in the dark to those tested in light (Fig. 2A).

**Figure 2.** Social preference requires visual observation of similarly aged fish. **A,** Schematic of the experiment to assess whether visual information is required for fish to show social preference. Following the acclimation period, three week old zebrafish are presented with a SC and monitored under both normal illumination and darkness, where the order of exposure to each condition was randomized. SPIs resulting from such experiments are indicated below schematics. **B,** Polar histograms of body orientations of the observer when on the SC side of the chamber during both light and dark sessions. The preference for the observer fish to orient at 45 degrees to the SC chamber is not present in darkness (thin lines indicate 2*SEM, n = 90), supporting the inference that such orientation in the light represents monocular viewing of conspecifics. **C,** Histograms of SPIs for all individuals during the dark and light conditions in the presence of a single fish as SC. **D,** Histograms of the SPIs of one week old fish observing three week old fish (left), and the SPIs of three week old fish observing one week old fish (right). See Supp. C for the SPIs in the absence of SC. Numbers in brackets indicate Mean SPI.

Removal of background illumination completely abolished the tendency of observer fish to orient towards the conspecific viewing chamber (Fig. 2B) and social preference was abolished, as evidenced by the distribution of SPIs (Fig. 2C, and Supp. Fig. 2A; Ac vs SC, p=0.57). Furthermore, with normal background illumination, replacing the transparent window with an opaque barrier also eliminated preference for the SC (Supp. Fig. 2B). These experiments provide strong evidence that the social preference behaviour of three week old zebrafish depends on vision.

The data above indicates that during the first three weeks of their life, zebrafish develop a robust social preference to view age-matched conspecifics. However, during this time they also change significantly in size, doubling their head to tail length (Supp. Fig. 1A). To assay whether the size/age of the SC fish influences social preference, we monitored the behaviour of one and three week old zebrafish presented with larger/older or smaller/younger conspecifics as the SC (Fig. 2D and Supp. Fig 2C).

One week old fish not only showed no preference for three week old fish, but also a slight aversion to viewing the larger fish, supporting the conclusion that the development of social preference reflects maturation of the observer and is not simply dependent on the age/size of the stimulus (Fig. 2D left; Ac vs SC, p=0.002; SPI difference (Ac-SC) = −0.111). Three week old fish also did not exhibit a strong social preference when presented with one week old fish as the SC (Fig. 2D, right; Ac vs SC, p=0.02; SPI difference (Ac-SC) = 0.106). However, the broader distribution of SPIs suggests that the smaller/younger fish may still influence the behaviour of the larger/older observing fish, which could be due to fish becoming progressively more responsive to any moving objects within their environment.

### Zebrafish coordinate their movement

Three week old fish display robust visually-driven social preference; high-speed videography additionally allowed us to investigate the extent to which the behaviour of the SC fish influenced the behaviour of the observer (Supp. Movie 5).

Young zebrafish tend to move in small bouts of activity consisting of discrete swims or turns (Orger et al., 2008). Individual bouts were detected by identifying a peak in the motion tracking signal (Fig. 3A, top trace). Averaging all of these bout time-courses (Fig. 3B) revealed a pre- and post-bout quiescent period, the timings of which reflected the periodicity of movement. These quiescent periods shortened from one to three weeks of age as the mean bout frequency increased ~50%, from 0.79 Hz at one week to 1.22 Hz at three weeks. As observed in other behavioural contexts, these movement bouts were composed of a mixture of forward swims and orienting turns (Fig. 3C)(Orger et al., 2008).

**Figure 3.** Development of the dynamics of social interaction. **A,** Example of the motion bout detection and alignment analysis: the left schematic shows the test chamber indicating in red the side in which the SC fish is visible and in blue the side in which it is not. The plots on the right indicate how movement bouts were analysed. Top plot shows movements bouts of the SC fish. Peaks in movement trajectories were identified with a dual-threshold algorithm (upper threshold dotted line is 3*standard deviation (3*SD) and lower threshold dotted line is 2*SD from baseline). The middle plot shows the movement bouts of the observer, test fish. The movement peaks of the SC fish were used to extract short time windows of the movement trajectories of the observer fish trajectory (2 seconds either side of the SC fish movement peak). The bottom plot shows the ‘bout-triggered-average’ (BTA) movement for the observer fish which was computed by averaging movements across all of the four second time windows aligned to the SC peak movement. BTAs were computed separately depending on whether the observer fish could view the SC or not (left schematic). **B,** The average bout motion time-course for all SC fish, normalized to the peak movement of each fish, at different developmental stages. The average bouts are overlaid to highlight changes in the kinetics between one and three weeks of age. **C,** Scatter plot presentation of all bouts, where each bout is represented by a single point that indicates the change (A) in position and orientation that occurred during that bout (n = 247779 bouts). **D,** BTAs of one- to three week old observer fish motion aligned to movement bouts of single SC fish (red) or plotted when the SC was not visible (blue) (one week: n = 106, two week: n = 136, three week: n = 163). **E,** BTAs for fish monitored in darkness when on the same (red) or opposite side (blue) of the SC (n = 90).

We next asked whether a motion bout produced by the SC fish influenced the movement of the observer. Short time windows of the motion trajectories from the observer fish, normalized by each individual’s average motion peak, were extracted and aligned to the bouts of the SC fish (Fig. 3A, middle trace) and were then averaged over all bouts (Fig. 3A, bottom trace). This generated a ‘bout triggered average’ (BTA), which is an estimate of how the motion bout of the SC influences movement of the observer.

A clear interaction between the movement of the SC fish and the observer was present at all stages of development. Notably, a bout of motion by a SC fish was, on average, coincident with a synchronous increase in motion by the observer fish. The strength of this motion coupling increased substantially over development (Fig. 3d), correlating with the enhancement in positive social preference. This visual coupling behaviour was, unsurprisingly, absent for fish in the dark (Fig. 3E). These results indicate that not only do three week old fish prefer to be with conspecifics, but that their behaviour is more strongly coupled with that of their social partners.

### Social preference and interaction are differentially impaired by drug exposure

We next assayed whether pharmacological manipulations that affect sociality in adult animals similarly influence the manifestation of social behaviour in young zebrafish.

Social learning in adult zebrafish is dependent upon N-methyl-D-Aspartate (NMDA) Receptor signalling(Maaswinkel et al., 2013) and we first assessed whether manipulating this pathway altered the social preference and interaction behaviour of zebrafish larvae. The NMDA receptor antagonist MK-801 was acutely administered at a concentration of 100 μM for 1 hour prior to assaying three week old zebrafish (Fig. 4A-D). Although MK-801 can lead to increased locomotor activity (Menezes et al., 2015), at this concentration and age treated zebrafish showed no overt change in the overall amount of swimming. However, we found that three week old fish exhibited no social preference (Fig. 4A and Supp. Fig. 2D; Ac versus SC period, p=0.51). This result suggests that blocking NMDA receptors interferes with circuitry required for social interactions, both in larvae and in adult fish. In addition, video-tracking revealed a significant alteration of movement dynamics in MK-801 treated larvae. The treated fish produced swim bouts lacking the pre- and post-bout quiescent periods (Fig. 4B), and a near total loss of conventional forward swims (Fig. 4C). Every bout involved a change in orientation (i.e. turning), which is consistent with the “erratic movements” observed in MK-801 treated adult fish(Sison and Gerlai, 2011) and reveals a substantial disruption to the movement control system after drug treatment. These altered bout dynamics also produced an asymmetry in the pattern of social interaction (Fig. 4D); the motion of the observer fish strongly influenced the movement of the untreated SC fish (dip prior to 0 ms), but the drug-treated observer was much less influenced by the movement of the SC fish (smaller dip after 0 ms; Fig. 4D). Furthermore, the synchronicity peak at 0 milliseconds lag was abolished.

**Figure 4.** Exposure to NMDA receptor antagonist or ethanol disrupts social preference and differentially impairs social interactions. **A**-**D**) Analysis of fish treated with 100 μM MK-801 NMDA receptor antagonist. **A,** Histogram of SPIs revealing no apparent preference for the SC and (inset) body orientations showed little or no direction towards the SC chamber (zero position). SPIs during the acclimation periods are shown in Supp. D. **B,** Average motion bout profile for MK-801 treated fish. Relative to untreated controls (grey plot), there is a reduction in the pre- and post-bout quiescent periods and consequently the periodicity of bout generation. **C,** Scatter plot presentation of all bouts (n = 85275 bouts) from all tested fish, where each bout is represented by single point based on the position and body orientation change that occurred for that bout. MK-801 treatment results in a conspicuous reduction in forward swimming bouts (‘0’ position on X-axis). **D,** Bout-triggered averages (BTA) of MK-801-treated observer fish when the SC fish was visible (red plot) or not (blue plot). There is a disruption of normal movement interactions before and after 0 seconds offset (compare to equivalent plots in or in h and l below) and the abolishment of behavioural synchrony at 0 seconds offset (see results text for further explanation). **E**-**H)** Comparable analyses as in A-D of fish treated with 0.125% alcohol. **E,** Plot of SPIs showing that social preference (red) remains and (inset) body orientations were directed towards the SC chamber. **F**-**H**, Average motion bout profiles (F), bout distributions (n = 101167 bouts) (G) and BTA plots (H) are all similar to untreated three week old zebrafish. **I-L**) Comparable analyses as in A-D of fish treated with 0.5% alcohol. **I,** Analysis of SPIs showing social preference is severely disrupted and (inset) body orientations are less strongly directed towards the SC chamber. **J-L,** Average motion bout profiles (J), bout distributions (n = 69675 bouts) (K) and BTA plots (L) are all similar to untreated three week old zebrafish.

Acute exposure to high concentrations of ethanol are also known to influence the social behaviour of adult zebrafish (Gerlai et al., 2000; Ladu et al., 2014). Consequently, we exposed three week old fish to low (0.125%) and high (0.5%) levels of ethanol 1 hour prior to and during testing in the social assay (Fig. 4E-L). The influence of ethanol exposure was concentration dependent. Fish exposed to low ethanol retained a strong SPI (Fig. 4E, and Supp. Fig. 2D; Ac vs SC period p=5.8*10^-8^) and their bout dynamics (Fig. 4F) and composition (swims vs turns) (Fig. 4G) were unaffected. Furthermore, the strength of their BTA interaction was similar to age-equivalent untreated fish. In contrast, upon exposure to a higher concentration of ethanol, social preference was greatly reduced and the SPI distribution was not significantly different from the acclimation period (Fig. 4L and Supp. Fig. 2D, Ac vs SC p=0.07). Remarkably, despite this loss of social preference by zebrafish exposed to high ethanol concentrations, movement dynamics (Fig. 4J), distributions of swim turns (Fig. 4K), and the strength of movement coupling with other fish (Fig. 4L) were not substantially affected. These intriguing results suggest that social preference and interactions with other individuals, each a fundamental component of social behaviour, can be decoupled by pharmacological, and likely other, manipulations.

## Discussion

### The development of social preference

We have shown that zebrafish gradually develop a “social” preference, which we define as the tendency to remain in a chamber that provides visual access to conspecifics; a behaviour that is absent in one week old fish, begins to emerge by two weeks, and is very robust at three weeks. This preference is visually driven and does not solely depend on the age/size of the conspecific partners as one week old larvae show no interest in larger three week old fish. These results suggest that social preference arises with the development of neural systems that appear or mature during the second and third weeks of life. For instance, it is known that some brain areas, such as the pallium (Dirian et al., 2014), undergo extensive growth during the establishing period of social preference.

Whether the preference to observe conspecifics reflects a drive to shoal/school or aggression(Gerlai et al., 2000) is difficult to distinguish with this assay. Both of these behaviours involve multi-modal stimuli (olfactory and tactile), which are prevented in our assay by the transparent barrier dividing the observer and social cue. Nevertheless, our assay demonstrates that visual cues alone can drive social behaviours and is thus easily translated to experimental setups that require restrained fish, but allow for detailed investigation of the underlying neural activity, e.g. two-photon microscopy.

We have not yet fully characterized the specific visual features that drive social preference. However, a simple preference for moving stimuli is unlikely to explain the response since three week old fish show little interest in viewing moving younger/smaller conspecifics. Our assay has demonstrated that visual stimulation is sufficient to drive social behaviour at three weeks of age. The presentation of visual “social” stimuli to restrained fish is much more straightforward than attempting to recapitulate the complex tactile and olfactory stimuli that are also involved in schooling/shoaling interactions (Cornelia H et al. 2012). Therefore, this social behaviour assay will considerably facilitate future studies to characterize the changes in neural circuitry that correlate with this fundamental behavioural transition.

It is intriguing that not all fish develop a positive response to conspecifics as some individuals exhibit avoidance behaviour when other fish come into view. This result warrants further investigation. For instance, our assay could be used to determine whether fish exhibiting aversive behaviour retain this negative social bias after multiple presentations of the SC or whether different environmental, pharmacological and genetic manipulations predispose developing zebrafish to express more aversive or attractive social behaviours.

### Social interaction as a coupled dynamic system

We found that when a fish observes the movement of a conspecific, its own swimming is affected. This visually-mediated coupling of movement is already present in one week old larvae, but strengthens considerably over the following weeks. Coupling the motion of one fish to that of another is an important prerequisite for the coordinated behaviour that predominates in groups of schooling fish (Miller and Gerlai, 2012). The temporal profile of this movement coupling, notably its remarkable synchronicity, is reminiscent of the coordinated movements apparent in other social organisms, including humans (Richardson et al., 2007; Sebanz et al., 2006).

However, synchronization of behaviour is observed in many physical systems with coupled dynamics, such as two metronomes on a shared surface (Pantaleone, 2002) as well as for many biological rhythms (Winfree, 1967). If any two periodic movement generators are sensitive to the motion of one another, then they will act as coupled oscillators and will have a natural tendency to synchronize. We have found such a coupling of observation of movement to movement generation in young zebrafish, but whether such coupling dynamics are important for the shoaling/schooling interactions of adult fish, or any other species demonstrating coordinated synchronous movements, warrants further investigation.

Disruptions to the coordination of behaviours, such as the loss of synchronized eye-blinking in autistic subjects (Sears et al., 1994; Senju et al., 2007), are now being identified as potentially important biomarkers of disease that may facilitate early diagnosis and intervention.

### Pharmacological manipulation of early social behaviour

The NMDA receptor antagonist MK-801 disrupted both social preference and interaction, whereas alcohol exposure disrupted only preference, leaving intact the ability of fish to couple their movements. This suggests that these two aspects of the social behaviour can be at least partially disassociated.

In addition to confirming that acute treatment with 100 μM MK-801 disrupts social preference in larvae, as was previously shown in adults(Sison and Gerlai, 2011), we also found that it greatly alters underlying movement dynamics. MK-801 treated larvae show strongly reduced bout periodicity and do not produce conventional forward swims, which could explain the observed deficit in coordinated behaviour. Such movement impairments will also affect the ability of treated fish to shoal, and might explain why adult zebrafish exposed to a lower concentration (5 μM) of MK- 801 exhibited disrupted shoal cohesion(Maaswinkel et al., 2013).

In contrast to NMDA receptor blockade, fish exposed to high concentrations of ethanol exhibited no disruption of intrinsic movement dynamics and show wild-type levels of movement interaction, but social preference was severely disrupted. These results highlight our assay’s sensitivity to distinguish components of social behaviour, preference and interaction, which could be separately impaired by different pathologies. Consequently this assay should be well suited for analysis of a range of genetic (Pietri et al., 2013) and pharmacological (Scerbina et al., 2012) manipulations that have been linked to developmental disorders affecting social behaviour.

### Future Directions

If we are to identify and characterize the neural circuits that underlie the development of social behaviour in zebrafish, it will be necessary to adapt our assay to enable the monitoring and manipulation of neural activity *in vivo* during social interactions. Fortunately, the brains of zebrafish are still small and relatively transparent during the developmental stages at which we observe the onset of social preference and coupled interactions, and are thus amenable to optical and optogenetic techniques for anatomical and functional investigation of whole-brain circuitry(Ahrens and Engert, 2015). Leveraging the power of these optical and genetic tools in young zebrafish, detailed comparison of the neural circuits established during normal and atypical development is likely to produce fundamental insights into the neural basis of social behaviour and its associated pathologies.

## Acknowledgments

*We thank Isaac Bianco, Jason Rihel, David Tingley and David Schoppik for helpful advice and comments on the manuscript, Carole Wilson, Heather Callaway, Karen Dunford, Jenna Hakkesteeg, and Matthew Wicks for fish care. This work was supported by Wellcome Trust Funding to S.W.W. and E.D. The authors declare no competing financial interests.*

## Author contributions

*ED, SWW, and ARK designed the project; ED performed experiments; GL created the software to acquire and analyze the data; ED and ARK analyzed data; ED, ARK and SWW wrote the paper.*

*The authors declare no competing financial interests.*

